# Automated Parallel Pattern Search Optimisation of Microfluidic Geometry for Extracellular Vesicle Liquid Biopsies

**DOI:** 10.1101/2021.11.13.468499

**Authors:** Colin L. Hisey, Arvin Lim, A.J. Tyler, Larry W. Chamley, Cherie Blenkiron, Richard J. Clarke

## Abstract

Microfluidic liquid biopsies using affinity-based capture of extracellular vesicles (EVs) have demonstrated great potential for providing rapid disease diagnosis and monitoring. However, little effort has been devoted to optimising the geometry of the microfluidic channels for maximum EV capture due to the inherent challenges of physically testing many geometric designs. To address this, we developed an automated parallel pattern search (PPS) optimiser by combining a Python optimiser, COMSOL Multiphysics, and high performance computing. This unique approach was applied to a triangular micropillar array geometry by parameterising repeating unit cells, making several assumptions, and optimising for maximum particle capture efficiency. We successfully optimised the triangular pillar arrays and surprisingly found that simply maximising the total number of pillars and effective surface area did not result in maximum EV capture, as devices with slightly larger pillars and more spacing between pillars allowed contact with slower moving EVs that followed the pillar contours more closely. We then experimentally validated this finding using bioreactor-produced EVs in the best and worst channel designs that were functionalised with an antibody against CD63. Captured EVs were quantified using a fluorescent plate reader, followed by an established elution method and nanoparticle tracking analysis. These results demonstrate the power of automated microfluidic geometry optimisations for EV liquid biopsies and will support further development of this rapidly growing field.

## 1 Introduction

Extracellular vesicles (EVs) are membrane-bound micro and nanoparticles that are extruded by all cells and circulate in all bodily fluids. Due to their immense potential as liquid biopsy targets, many affinity-based microfluidic technologies have been developed to isolate EVs of certain subtypes from patient or cell culture samples^1–3^. As with circulating tumour cell (CTC) chips, microfluidic capture of EVs often involves the functionalisation of antibodies or other capture ligands on the inner surfaces of microchannels. Upon contact, depending on binding affinity, shear forces, and other factors, there is a probability that EVs will be captured for either on-chip analysis, lysis, or elution^4–7^.

This probability of surface capture is highly dependent on the internal geometric structures, and existing microfluidic liquid biopsy devices vary significantly, from simpler designs like the micropost arrays^8–11^, alternating shrinking and expansion of channel widths^5^, and staggered herringbone grooves^4,6,7^, to more complex devices utilising multiple elements, such as “NanoVelcro”^12^. These geometries are required because the highly laminar nature (low Reynolds number) of the fluid flow within the channels limits any mixing to diffusion, which is ineffective if maximum surface interactions are desired. Thus, the established geometries increase the surface interaction probability by either greatly increasing the total surface area or disrupting the highly linear flow profiles present in straight channels.

Pillar arrays were first used to increase the total area of antibody-functionalised surfaces. Their geometry has been somewhat optimised for capturing circulating tumour cells (CTCs), which are two orders of magnitude larger than small EVs. However, micropillar designs have been shown to have a limiting capture efficiency of 65 percent for CTCs due to their inherent laminar flow^9^. For EV capture, ciliated micropillar arrays have been used in a sieve-based filtration device^11^, and only one study has demonstrated EV immunocapture using Y-shaped microposts and a unique nanostructured graphene surface^13^. More complex structures, including staggered herringbone grooves, have been used to induce microvortices within the flow, effectively increasing interactions between the particles and the surface of the channel^4,14,15^. These geometries were initially designed within the context of microfluidic mixing^16^, but have more recently been optimised for particle-surface interactions for immunoaffinity capture systems^14,17^. Another approach, named “ExoChip”, involves rapid shrinking and expansion of channel diameters to enhance fluid mixing in thinner channel sections and slowing the flow rate of EVs in the larger regions^5^.

To assess the performance of microfluidic devices, numerical models are often used in favour of laboratory testing due to their adaptability, ease of construction, and low cost. These models solve the Navier-Stokes equations, which govern flow of a Newtonian fluid, and usually make several simplifying assumptions. Although simple micropillar channels have analytical solutions for the average longitudinal velocity^18^, numerical simulations are advantageous, despite their larger computational complexity, due to their ability to describe the full velocity field. This enables the prediction of the advection of suspended particles in flow, which is necessary to accurately quantify the particle capture. Both micropillar and staggered herringbone designs have been optimised to some degree using computational fluid dynamics (CFD) simulations using commercially available software. CFD solves the Navier-Stokes equations in a top-down manner by discretising the continuum equations using a finite-element method^19^. However, to the best of our knowledge, no microfluidic geometries have been computationally optimised in the context of EV immunoaffinity isolation.

In one study, a CTC-capture micropillar geometry optimisation study was conducted for several parameters including a qualitative array type of either square, diagonal or triangular pillars^9^. In another study, grid-search optimisation was conducted for a staggered herringbone groove design for CTC-capture using both channel and groove depth, width and height, number of grooves per half cycle, groove angle, groove pitch and an asymmetry factor^14^. To quantify the frequency of particle/surface interactions this study employed point particles with specified virtual radius, that were tracked and said to have a chance of capture if the distance between the particle and the surface became less than the radius. This virtual radius was initially developed in 2012 for micromixing applications^17^, and it was later used in conjunction with lattice Boltzmann methods, which is a bottom-up approach to solving the Navier-Stokes equations^20^. This approach is in contrast to the usual top-down process of discretising the equations. Due to its natural divisibility, the bottom-up approach is better suited to computational parallelisation^21^. In another study, staggered herringbone grooves were used in a surface-particle optimisation using a modified Arrhenius model to describe the particle binding reaction, which better describes the chemistry between the suspended particles and the capture ligands on the surface of the channel^22^.

We report the development of an automated parallel pattern search optimiser for improving the design and effectiveness of microfluidic EV capture devices. We outline the justification of several assumptions, the unexpected results for maximum particle capture efficiency, and experimental validation of the findings by fabricating channels and testing them using bioreactor-produced breast cancer cell line-derived EVs. As the field of microfluidic EV liquid biopsies continues to grow, this type of automated geometry optimisation could be invaluable in enabling researchers to more efficiently capture EVs of interest without spending extensive amounts of time physically optimising their device designs.

## 2 Results and Discussion

### 2.1 Optimised Designs

The optimisation workflow was initialised using a microchannel geometry consisting of a triangular arrangement of circular pillars of radius *r* = 30*µ*m and separation parameter *a* = 3.5 (where centre-to-centre pillar separations distances *d* are given by *d* = *ar*). After 25 objective evaluations the solution converged to an optimal design with *r* = 20*µ*m and a = 3.56. The objective function (capture rate) under this design was evaluated as 391, a 140% increase in performance compared to the starting geometric design. The optimisation was repeated for three different sets of initialisations of *r* and *a*. The resulting objective functions can be found in Figure 1 and Table 1.

**Fig. 1.**
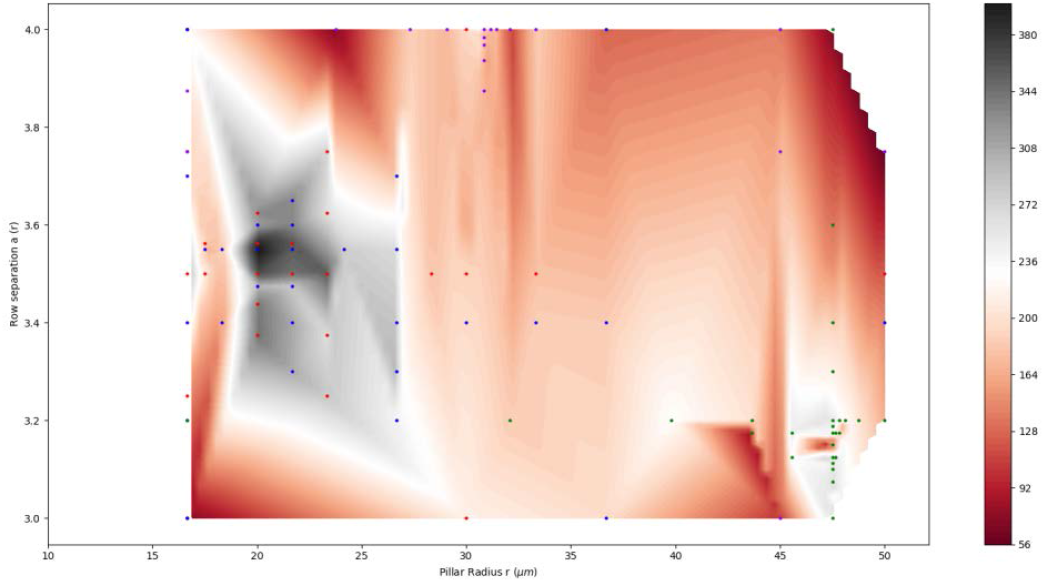
Predicted EV capture rate plotted as a contour map against geometric design parameters *r* and *a*, for four instances of the PPS optimisation solver. The dots represent all solutions evaluated with their different colours indicating different solver initialisations, while the contour gradient of black to burgandy indicate the particle capture efficiency from highest to lowest, respectively.

**Table 1.** Comparison of initial and optimised geometries, as computed by the PPS algorithm. Subscripts I and O denote initial and optimised values.

| $r_I$ | $a_I$ | $EV_I$ | $r_O$ | $a_O$ | $EV_O$ | % change |
| --- | --- | --- | --- | --- | --- | --- |
| $3 \times 10^{-5}$ | 3.50 | 163 | $2 \times 10^{-5}$ | 3.5625 | 391 | 139.88% |
| $4.75 \times 10^{-5}$ | 3.20 | 264 | $4.73 \times 10^{-5}$ | 3.125 | 301 | 14.02% |
| $4.5 \times 10^{-5}$ | 3.75 | 164 | $3.08 \times 10^{-5}$ | 4 | 224 | 36.59% |
| $3.66 \times 10^{-5}$ | 3.4 | 184 | $2.00 \times 10^{-5}$ | 3.55 | 405 | 120.11% |

It can be seen that the initial starting point affects the performance of the optimisation, with two of the four initialisations converging to a geometry with lower EV capture rate. This is to be expected, since the nonlinearity of the problem necessitates the use of a heuristic optimisation routine (like the Parallel Pattern Search algorithm used here), which only guarantees convergence to a local optimum. Hence the need for a number of initialisations for a good survey of geometry design space.

In order to highlight the difference between the geometry design with the highest and lowest EV capture rates, Figure 2 shows plotted flow streamlines in those microchannels. The best design had a capture rate of 405, whereas the worst design (found at any point during the optimisation routine) had a capture rate of 69. The better design had a less dense packing of pillars, as evidenced by both fewer pillars across the width of the microchannel and a shorter length of the periodic unit (although the total area of the microchannels in both cases is the same). This was a somewhat surprising finding, since a denser packing leads to a greater overall functionalised surface area. However, we see in the higher performing, lower density pillar arrangement that the streamlines follow the contours of the pillars more closely, providing greater opportunity for EV capture.

**Fig. 2.**
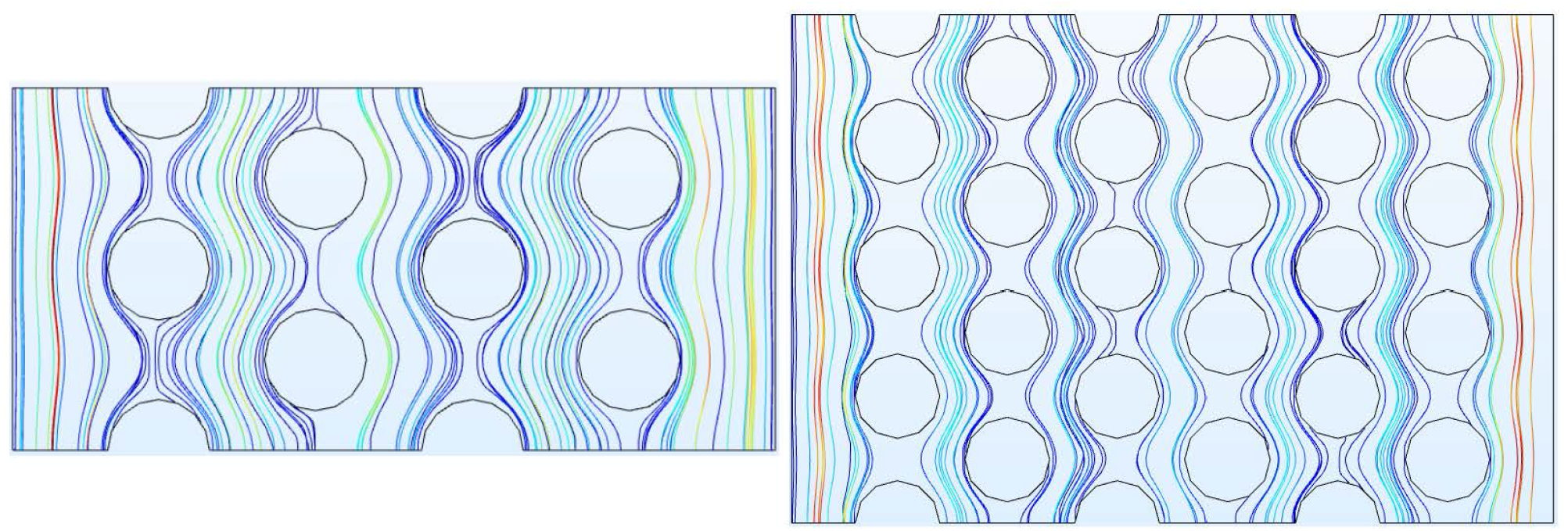
Streamlines for (left) best design (*r* = 20 : 0*µ*m, *a* = 3.55) and (right) worst design (*r* = 16.7*µ*m, *a* = 3.00) (right) Red streamlines correspond to higher velocities

### 2.2 Experimental Validation

To experimentally validate the designs shown in Figure 2, devices were fabricated using standard photolithography and soft lithography methods. Given the relatively high aspect ratio features, scanning electron microscopy (SEM) and optical profilometry were used to ensure that the pillars were not destroyed during the demoulding process. As shown in Figure 3, SEM images and optical profilometry demonstrate that the desired geometries were successfully produced in the final microchannel structural features (*r* = 20*µ*m, *a* = 3.55 and *r* = 16.7*µ*m, *a* = 3.00) throughout the entire length of the channels.

**Fig. 3.**
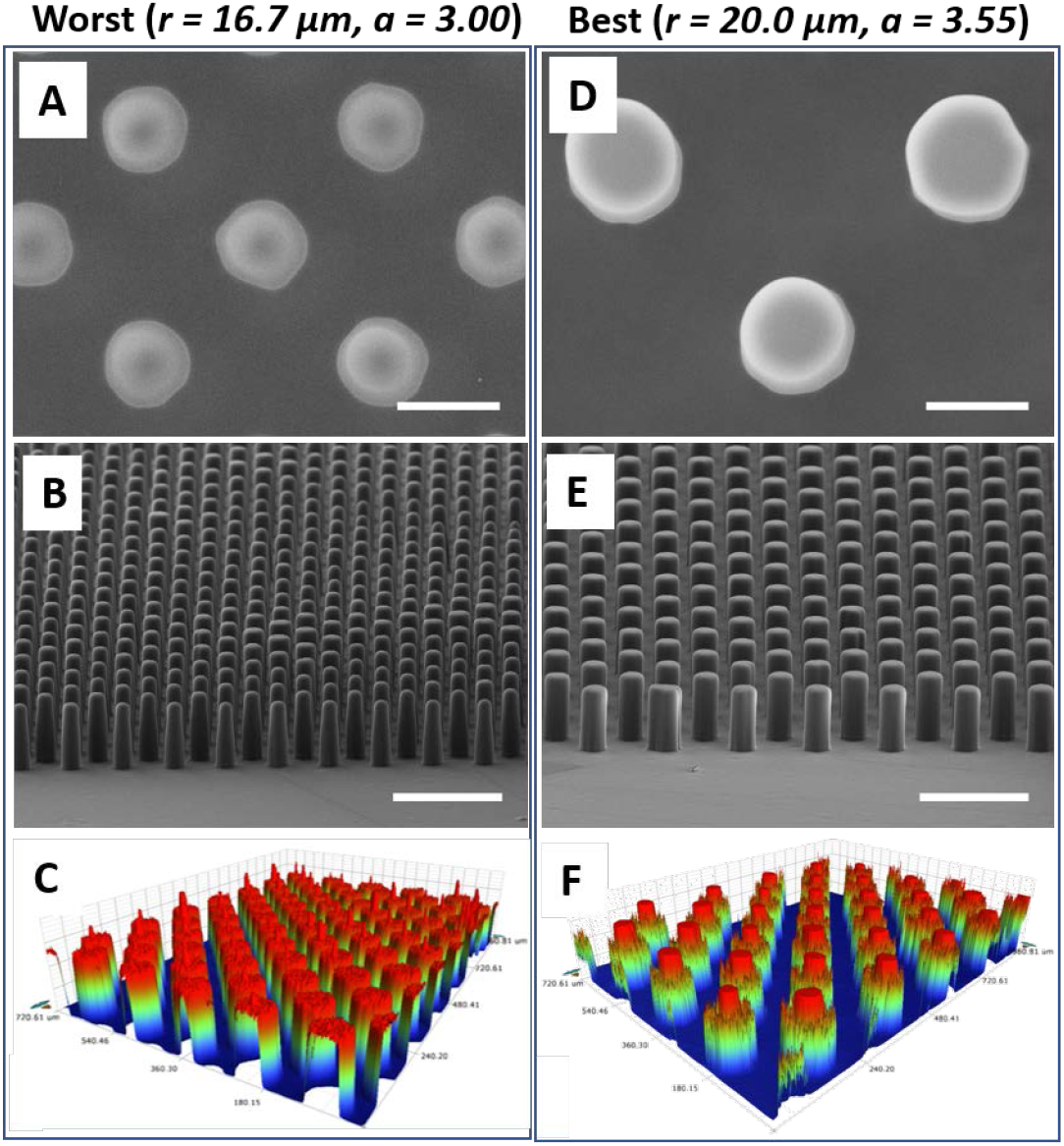
SEM and profilometry of the worst (A-C) and best (D-F) microfluidic channel designs as determined by the PPS optimiser. (A,D scale bar = 40 *µ*m, B,E scale bars = 150 *µ*m)

Two complimentary approaches were then tested for their ability to quantify the relative EV capture efficiencies between the two designs. In the first, fluorescently labeled EVs that were collected from BT-20 cells cultured in a bioreactor were quantified on-chip by placing the chips with captured EVs into a custom 3D printed plate reader frame. After correcting for the autofluorescence resulting from the chip structure and antibody functionalisation surface chemistry, the worst design from the simulations produced an average fluorescence signal of 301012 *±* 37925 a.u., while the best design produced a fluorescence signal of 607795 *±* 96760 a.u. as shown in Figure 4*A*. This represents an increase in capture efficiency of 102%, which is much less than the 6-fold increase expected based on the simulation results, but does validate there is an increase in EV capture efficiency between the worst and best designs.

**Fig. 4.**
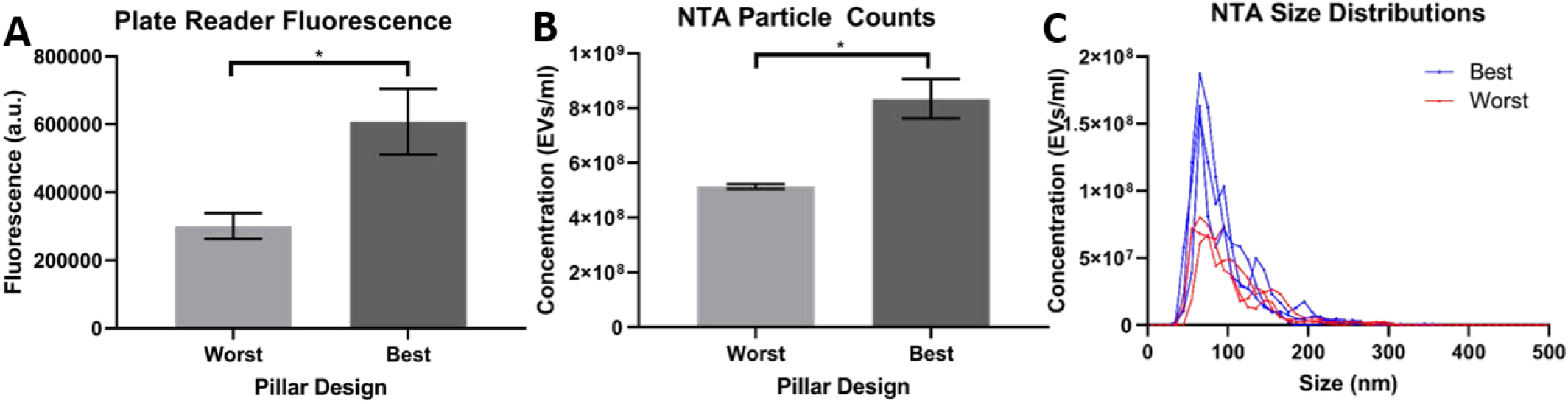
Comparison of EV capture quantification methods showing (A) normalised fluorescence signals from plate reader, (B) NTA particle counts following elution of captured EVs, and (C) NTA size distributions of eluted EVs. (n=3,^*^=p<.05)

**Fig. 5.**
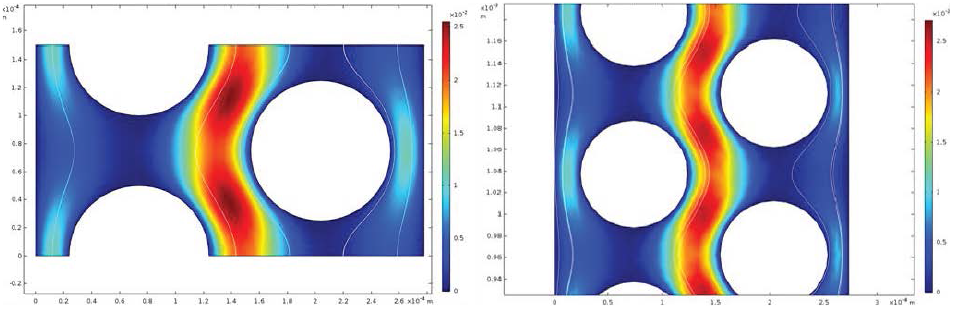
Flow around triangular pillar arrays simulated as assuming a periodic repeating unit (left) and explicit full geometry (right). Both designs comparisons show near-identical flow fields.

Assuming all the normalised fluorescence signal was from EVs, the plate reader approach agreed, to some degree, with the simulation. However, it is not a well-established technique, so it was compared to a single particle counting NTA method by using an affinity chromatography system where the EVs are released from the channel surfaces using acidic buffer and immediately neutralised downstream. This process was performed using the exact same devices that were quantified using the plate reader, and NTA results again demonstrated an increase in EV capture between the worst and best designs, from 5.14 × 10^8^ *±* 9.07 × 10^6^ EVs/ml in the worst design to 8.34 × 10^8^ *±* 7.19 EVs/ml in the best as shown in Figure 4*B*. This represents a 62% increase in capture efficiency. In addition, the size distributions of EVs eluted from the different channel geometries appear to be similar and again illustrate the differences in concentration, shown in Figure 4*C*. Although there are clear differences between these two EV quantification approaches, they collectively demonstrate that the simulations can be used to improve EV capture efficiency in microfluidic liquid biopsy applications.

The differences seen between the two EV quantification methods could be due to several reasons, including incomplete elution of the EVs from the capture surface or even elution at different rates given the different geometries. In addition, the fluorescent plate reader cannot provide a single particle count or any size distribution information, and the strong levels of background fluorescence could have resulted in imperfect normalisation of the total fluorescence signal. In the future, a standard curve could be created to give some indication of absolute EV quantity. This would also need to be done for each specific geometry, again due to the different levels of background fluorescence in each design. The differences seen between both experimental EV quantification methods and the simulations could be caused by several factors, such as the EV capture being a boolean function based solely on proximity in the simulations, whereas the imperfect kinetics of antibody binding to EVs mean the overall capture probability is inherently lower in the experiments. In addition, the simulations were performed using EVs with an assumed diameter of 100 nm. This is likely more in line with NTA-based quantification, which preferentially quantifies particles in this range due to its well-established decrease in accuracy when quantifying smaller particles^23^. Fluorescence quantification, on the other hand, would obtain a signal from EVs below the detection limit of NTA, and could in part explain the increased difference in EV capture measured from the plate reader analysis.

In the future, this simulation approach could be improved by relaxing many of the assumptions that were made in this study and expanding this approach to other device geometries. For instance, several assumed constants could instead be added to the explored parameter space, including the channel height, shape and angle of the pillar arrays, or flow rate. In addition, other common microfluidic geometries such as alternating staggered herringbone grooves could potentially be paramterised, optimised, and experimentally validated in a similar manner.

## 3 Conclusions

In this study, we demonstrate that a parallel pattern search optimiser built in Python could use COMSOL Multiphysics’ Particle Tracing Module to optimise microfluidic geometries for EV immunoaffinity capture. Surprisingly, the simulations suggested that simply maximising the total number of pillars, and thus the effective surface area, was not the optimum triangular pillar array geometry. Rather, slightly larger and more spaced out pillars allowed slower contact with EVs and encouraged them to follow the coutours of the pillars more closely. These findings were validated experimentally using fluorescently labeled EVs from a bioreactor and an antibody against the tetraspanin CD63, and quantified using a fluorescent plate reader as well as an acidic elution approach followed by NTA. These results demonstrate the potential of automated microfluidic optimisations for EV liquid biopsies and will support further development of this rapidly growing field.

## 4 Materials and Methods

### 4.1 Microchannel Design

We consider a microchannel of length 75mm, width 300*µ*m and height 100*µ*m. The EV suspension passed through the channel in PBS (phosphate buffered saline), and has a density of 1007 kg/m3 and a dynamic viscosity *µ* of 3.30 *×* 10^*-*3^ Pas. The volumetric flow rate (V) is 2.5 *×*10^*-*10^ m^3^/s. The pillar array within the microchannel is the same as^14^, an equilateral triangle arrangement of circular pillars with radius *r*, centre-to-centre (row) separation *d* = *ar* and corresponding lateral (column) separation 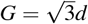. Observing manufacturing constraints, we explore the parameter range 50*/*3*µm < r <* 50*µm* and 3 *≤ a ≤* 4.

### 4.2 Design Optimisation

The modelling framework consists of several stages. For a given microchannel geometry, we simulate the flow of an EV laden fluid through that geometry, and determine the EV capture rate. The flow simualator is coupled to an optimisation routine that updates the microchannel geometry to maximise this capture rate. We provide more details below on these two components.

#### 4.2.1 Flow Simulations

At the small length scales found in microfluidic devices, viscous forces dominate inertial effects, and the flow can be well-described by the Stokes Flow equations (sometimes known as creeping Flow)^24^.

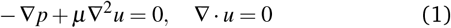

where *u* represents flow velocity, *p* flow pressure, and *µ* dynamics viscosity. We assume that the EVs have negligible inertia and are sufficiently dilute that they are advected by the flow without appreciably affecting it. As such, the trajectory of the EVs satisfy

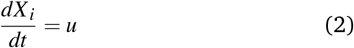

where *X*_*i*_ denotes the location of the *i*th EV in the suspension.

We assume that there is no flow on solid surfaces of the microchannel geometry. Moreover, due to the repeating nature of the geometric patterns within the microfluidic channel, the flow geometry can be reduced to a repeating unit cell with a periodic velocity boundary condition. This simplification provides drastic reductions in computational expense as only a small section of the entire device needs to be simulated. However, a pressure difference across the repeating unit cell is required, and with a difference of 1.108Pa found to give the require volumetric flow rate through the channel. To validate the periodic geometry approach, the flow through an entire microchannel, containing repeating arrangements of triangular or circular micropillars, were compared to the flow through a single repeating unit. As illustrated in Figure 1, simulations showed that the flow computed in the full geometry (right image) is seen to agree with that computed using a single periodic repeating unit (left image).

The flow equations and particle tracking and capture were undertaken using the Microfluidic and Particle Tracing modules within COMSOL Multiphysics (version 5.4). Every 0.1 seconds, for a period of 10 seconds, 100 particles were injected into the microchannel at randomly distributed locations across the channel inlet. An accumulator counter was created to track the number of particles (EVs) making contact with the wall (enabling particle capture), which was used as a measure of the design’s performance. A physics controlled unstructured tetrahedral mesh was used in simulations, with mesh convergence achieved using elements sizes in the range 6.71 *×* 10^*-*6^ to 2.24 *×* 10^*-*5^ (see Figure 6).

**Fig. 6.**
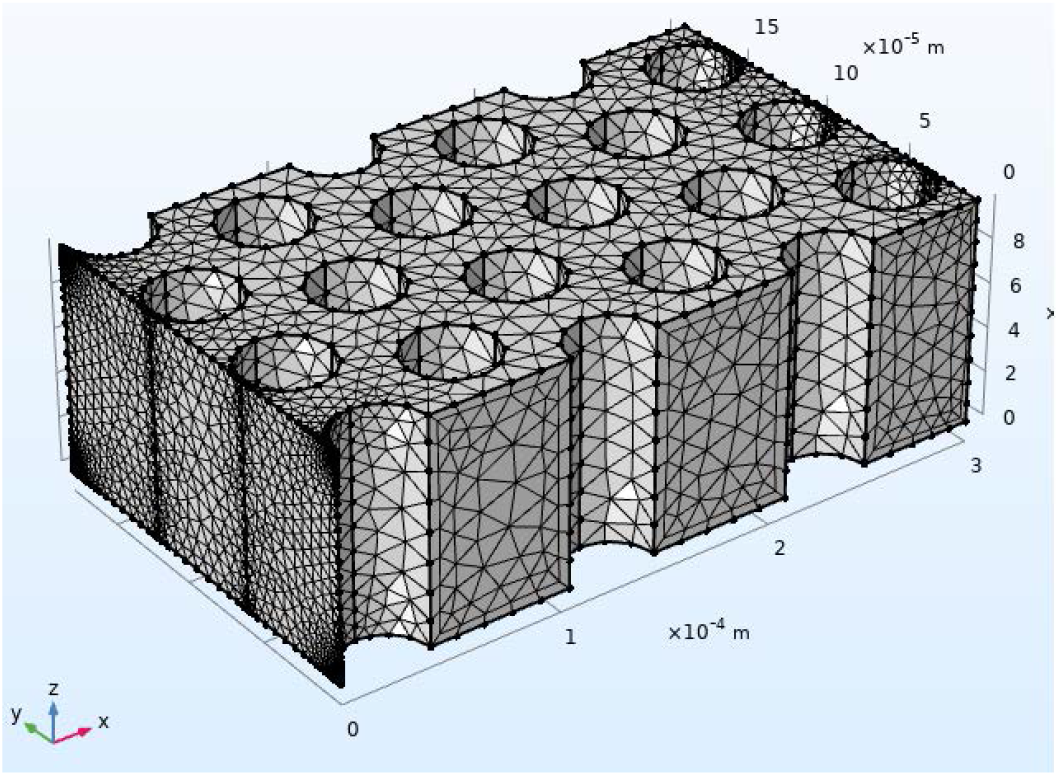
Computational mesh for periodic repeating unit of triangular array of circular pillars in a 300*µ*m wide microchannel.

#### 4.2.2 Parallel Pattern Search Optimisation

In this study, we implement a Parallel Pattern Search (PPS) optimiser to maximise the theoretical immunoaffinity-based capture of EVs on triangular micropillar arrays within a microfluidic channel. Parallel Pattern Search (PPS) is a gradient-free and parameter-based optimisation algorithm specifically designed for computationally expensive and bound-constrained problems of high dimensionality^25^. This algorithm starts at the outer edges of the parameter domain and works its way towards the centre. If an improved solution is found, the pattern shifts and continues converging. This algorithm is partly non-sequential, in that multiple points within the parameter domain can be simulated simultaneously, reducing the total computation time if high performance resources are available.

Our workflow combined a Python implementation of PPS coupled to the COMSOL particle tracing simulations. On each iteration of the PPS algorithm, geometric design parameters *r* and *a*, corresponding to pillar radius and spacing, are passed to the COMSOL scheme which simulates the flow and particle trajectories in the corresponding microchannel geometry, and returns to the PPS the particle capture for that particular geometric design. Convergence of the PPS algorithm was set to be when variations in the geometric parameters *r* and *a* become less than 5%, as these represent changes below manufacturing tolerances. All optimal designs were computed within 24 hours using 8 CPU’s and 16 GB of memory on the Mahuika New Zealand eScience Infrastructure (NeSI) high performance computing cluster.

## 5 Experimental Setup

### 5.1 EV Production and Isolation

EVs were produced using a two-chambered CELLine AD1000 bioreactor (Sigma) culture of BT-20 cells, a triple negative breast cancer cell line, as previously reported^26,27^ Briefly, cells were inoculated in complete media and and slowly adapted to serum-free advanced media with serum replacement CDM-HD (Fiber-cell). Once per week, the 500 ml media chamber was refreshed and twice per week, the 15ml of conditioned media in the cell chamber was collected and used for EV isolation. EVs were isolated by first centrifuging at 2,000 xg for 10 min to remove dead cells and large debris, followed by 10,000 xg for 30 min to pellet large EVs, and finally 100,000 xg for 70 min to pellet small EVs. Small EVs were then resuspended in 500 *µ*l PBS, labeled with 10 *µ*M Vybrant DiO (V22886, ThermoFisher), and purified using a 35 nm qEV Original size exclusion chromatography column and an automated fraction collector (Izon). EV-rich fractions (7-10), as validated previously based on nanoparticle tracking analysis (NTA) and bicinchoninic acid assay (BCA)^26,27^, were pooled and used for the remainder of the study.

### 5.2 Microfluidic Fabrication and Characterisation

Microfluidic devices were fabricated using standard SU8-2100 photolithography and PDMS moulding. First, 80 mm diameter acrylic wafers were cut using a laser cutter. SU8-2100 (MicroChem) was then spincoated and processed with appropriate baking, exposure, and development conditions. Wafers were then treated with HMDS (Sigma) overnight in a vacuum degasser to improve demoulding of the pillar features. A 10:1 ratio of Sylgard 184 PDMS (Dow Corning) was mixed, degassed, and poured over the patterned wafers, degassed again, and cured at 60 °C for 3 h. PDMS channels were then demoulded from the patterned wafer and connector holes were punched using a filed 18g needle. Channels and glass slides were plasma treated with air at 40W for 2 min (Harrick Plasma) and left for 10 min before introducing a series of linker chemistries. To validate the desired channel geometries, SEM imaging was performed by coating devices with 10 nm of gold (Q150R S, Quorum) then imaging using a JCM-6000 benchtop SEM (JEOL) at 15 kV. Optical profilometry was used to further validate the structures using a Contour GT-K Optical Profiler (Bruker).

### 5.3 EV Capture Efficiency Quantification

10 min after bonding, microfluidic channels (n = 3 of each design) were functionalised with an antibody against CD63 (SantaCruz, sc-5275) by using a well-established linker chemistry^4,28–30^. Briefly, 8% percent (3-Aminopropyl)triethoxysilane (APTES, Sigma) in ethanol was flowed at a rate of 20 *µ*l/min for 15 min then left stagnant for 20 min, followed by a flush with pure ethanol. Then, 4% glutaraldehyde in water was flowed and left stagnant similarly, with a similar water flush. Finally, 100 *µ*l of CD63 antibody (MX-49.129.5, sc-5275, SantaCruz) in PBS at a concentration of 5 *µ*g/ml was flowed through at a rate of *µ*l/min until all but roughly 5 *µ*l was left in the inlet tubing. The inlet tubing was then cut at the fluid mark, and the chips were stored at 4 °C overnight. The following day, the chips were flushed with PBS at 20 *µ*l/min for 15 min, then the BT-20 EVs were flowed through at a rate of 15 *µ*l/min. The chips were once again flushed with PBS at a rate of 15 *µ*l/min.

We tested whether a plate reader could be used to compare the capture efficiency of the DiO-labeled EVs in each chip design (Supp Fig 1). Using a custom 3D printed chip holder with alignment markers based on a 96-well ELISA plate (.stl file provided in supplementary), chips were loaded and fluorescence was quantified for 483 nm excitation and 501 nm emission on an Ensight Multimode Plate Reader (Perkin Elmer). Chips that were functionalised similarly but did not undergo EV capture were used as background fluorescence controls (antibody only). Immediately following fluorescence analysis, EVs were eluted with 200 *µ*l pH 2.2 Glycine-HCl buffer into 20 *µ*l pH 8.5 Tris-HCl neutralisation buffer as shown in Figure 7, then counted using NTA.

**Fig. 7.**
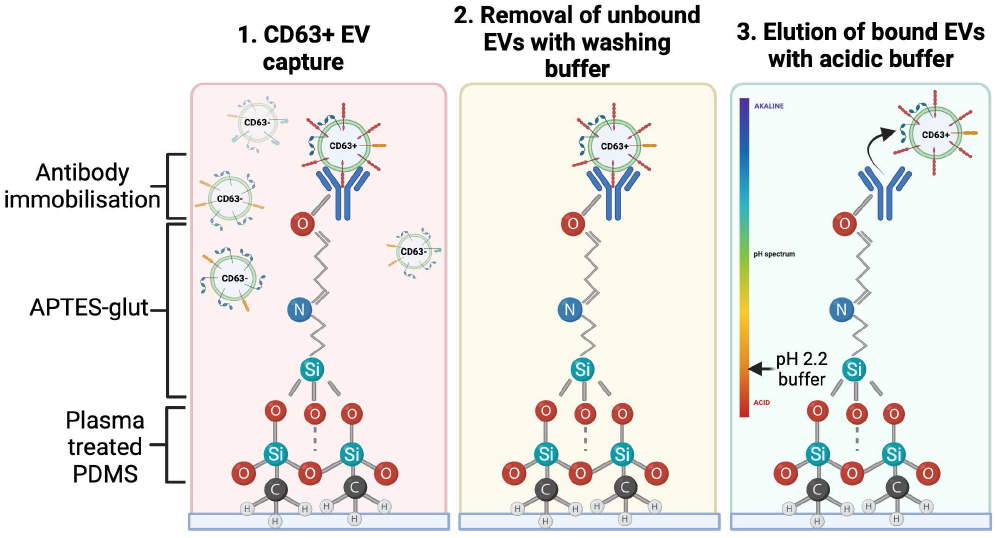
Schematic of EV capture and release approach.

### 5.4 Statistical Analyses

All averages are reported as average *±* standard error mean. Error in all figures are standard error mean. Statistical analyses were performed were unpaired t-tests (p<.05).

## Author Contributions

Colin Hisey: Conceptualization, Methodology, Validation, Formal analysis, Investigation, Resources, Writing – Original Draft and Reviewing Editing, Supervision, Project administration, Funding acquisition AJ Tyler: Methodology, Software, Validation, Data Curation, Visualization, Formal analysis, Investigation, Writing – Original Draft and Reviewing Editing Arvin Lim: Methodology, Software, Validation, Data Curation, Visualization, Formal analysis, Investigation, Writing – Original Draft and Reviewing Editing Lawrence W. Chamley: Resources, Writing – Review Editing Cherie Blenkiron: Resources, Writing – Review Editing Richard Clarke: Conceptualization, Methodology, Validation, Formal analysis, Investigation, Resources, Writing – Original Draft and Reviewing Editing, Supervision, Project administration.

## Acknowledgements

The authors would like to thank the Breast Cancer Foundation New Zealand Technology and Innovation Project Grant for funding this project, as well as the Hub for Extracellular Vesicle Investigations and New Zealand eScience Infrastructure (NeSI) for their continued support.

