## Supplementary figures and images for "Automated Parallel Pattern Search Optimisation of Microfluidic Geometry for Extracellular Vesicle Liquid Biopsies"

### Supp Fig 1

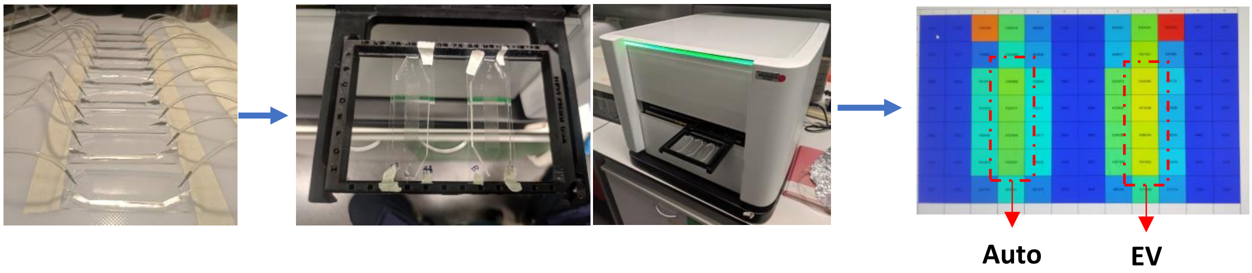


**Supplementary Figure 1.** Fluorescent plate reader EV quantification approach.
